# Development of a High-throughput Assay to Identify Novel Reactivators of Acetylcholinesterase

**DOI:** 10.1101/2025.07.16.664887

**Authors:** Becky L. Hood, Stefan Vasile, Gideon Shapiro, James D. Talton

**Affiliations:** Alchem Laboratories Corp., Alachua FL; Pharmorquest Inc.

## Abstract

Acetylcholinesterase (AChE) is a key enzyme in the nervous system capable of breaking down acetylcholine (ACh). ACh is a critical neurotransmitter that signals between nerve cells and muscles. Organophosphates (OPs), used as insecticides and nerve agents, irreversibly inhibit AChE by forming a covalent bond with the enzyme’s active site. Since exposure to these nerve agents can quickly lead to death, many researchers have strived to discover new drugs that will reverse the inhibition of AChE by such agents. The goal was to develop a high-throughput miniaturized assay for testing large numbers of compounds for AChE reactivation quickly and reliably. Here we describe our miniaturized assay that utilizes the sarin surrogate nitrophenyl isopropyl methylphosphonate (NIMP) in an Amplex Red coupled assay and its validation with 42 compounds selected from an *in silico* virtual screen.

## Introduction

Organophosphorus nerve agents such as isopropyl methylphosphonofluoridate (sarin) and S-2-diisopropylaminoethyl O-ethyl methylphosphonothioate (VX), are potent acetylcholinesterase inhibitors. Exposure to these agents can rapidly lead to the accumulation of the neurotransmitter acetylcholine in neuromuscular junctions and cholinergic synapses [1]. These reactive organophosphates inhibit the hydrolytic action of acetylcholinesterase (AChE) by phosphorylating the serine of AChE’s active site [2]. Phosphorylation is persistent because spontaneous reactivation is slow [3]. If left untreated, organophosphate (OP) poisoning leads to dose-dependent toxic signs including tremor, hyper secretion, seizures, respiratory depression and eventually death. Various quaternary oxime reactivators such as Pralidoxime (2-PAM) [4] and obidoxime [5], combined with anticholinergics such as atropine and scopolamine have been used to treat acute poisoning by OP nerve agents while seizures are often controlled by the administration of GABA receptor agonists such as diazepam or midazolam [6]. Organophosphorus nerve agents pose a special threat to the military: including active-duty combat personnel, nerve agent stockpile handlers, and nerve agent demilitarization workers [7].

Cholinesterase reactivators are generally used as antidotal therapies for pesticides and nerve agent poisoning [8]. The discovery of 2-PAM as a reactivator of OP inactivated AChE [9] led to numerous mono- and bis-quaternary oximes AChE reactivators with activity *in vitro* and *in vivo*. 2-PAM, obidoxime [10], and asoxime chloride (HI-6) [11] were fielded as OP nerve agent antidotes for combat soldiers around the world [12]. However, quaternary oximes are not able to cross the blood–brain barrier (BBB), thus limiting their therapeutic efficacy. Therefore, it is of great interest to identify novel reactivators that are brain permeable. To this goal, Amitai et. al. [13] discovered four novel reactivators of both sarin- and VX-inhibited butyrylcholinesterase (BChE) which also directly detoxified Sarin. These authors expect that their new reactivators will be permeable to the BBB. To date, compounds with broad spectrum reactivation are limited. Identifying classes of compounds to fill the gaps would be very beneficial.

The assay that is described here for identifying novel reactivators of OP inactivated AChE utilizes the sarin surrogate NIMP which was designed to phosphorylate cholinesterases with the same moiety as sarin, making it highly relevant for the study of cholinesterase reactivators [14, 15]. NIMP is a potent inhibitor of rat brain, skeletal muscle, diaphragm, and serum AChE as well as human erythrocyte AChE and serum BChE in vitro. Although safer to use than sarin and other nerve agents, keeping the exposure to a minimum is desired. Having a low volume assay using nanoliter quantities of NIMP was our goal for safety reasons as well as for cost savings.

For this assay we chose to use an Amplex Red fluorescent assay to measure the acetylcholinesterase activity. It is based on the coupled Amplex Red fluorescent assay available commercially from Invitrogen (brand of Fisher Scientific) [16, 17], which is more sensitive than the Ellman’s assay. Unlike the Ellman’s assay, compounds with sulfur do not interfere with the Amplex Red assay. An additional advantage of the Amplex Red assay is that it uses the natural substrate (acetylcholine) whereas the Ellman’s assay uses the acetylthiocholine. However, the Amplex Red assay is not a perfect assay because some fluorescent compounds or fluorescent quenchers will interfere with the fluorescent signal of the Amplex Red assay. The issue with fluorescent compounds appearing as reactivators is overcome by taking multiple time points. A fluorescent compound will not increase its fluorescence with time whereas a true reactivator will increase the fluorescent signal with time as the substrate is turned over by enzyme activity. Also, quenching (such as 12.5 µM HI-6, see Figure 6) will eventually be overcome by the enzyme activity at later time points. We see this in the dose response curves (DRCs) HI-6. The curves become well-behaved S curves at later time points in the assay. Figure 1 shows the coupled reaction involved in this assay.

**Figure 1:**
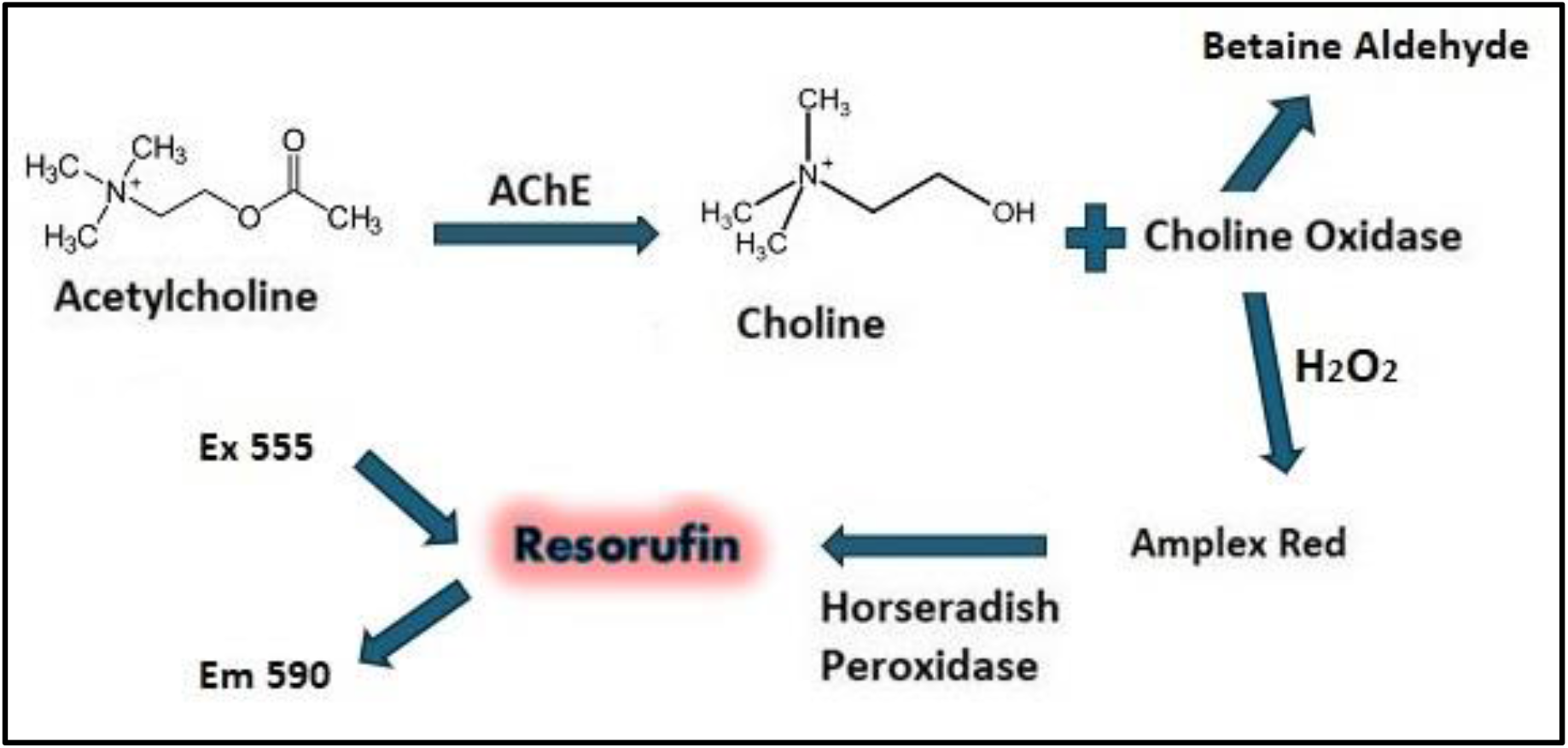
Amplex Red Assay Reaction. Acetylcholinesterase hydrolyzes acetylcholine to choline. Choline is then oxidized by choline oxidase, producing betaine aldehyde and H_2_O_2_. H_2_O_2_ reacts with Amplex Red, catalyzed by HRP, producing resorufin which can be quantified by fluorescence.

## Materials and Methods

### Reagents and Instruments

The sarin surrogate isopropyl (4-nitrophenyl) methylphosphonate (NIMP, cat # SY247222) was purchased from Acello Chembio (San Diego, Ca). Amplex Red reagent (cat # A12222) was purchased from Thermo Fisher Scientific (Waltham, Massachusetts). Horseradish peroxidase (cat # P8375), Choline Oxidase (cat # C5896), acetylcholine chloride (cat # A2661), human recombinant acetylcholinesterase (cat # C1682), HI-6 (cat # SML0024), 2-PAM (cat # 1553000), dimethyl sulfoxide (DMSO, cat # D2650) and Tris-HCl (cat # T3253) were purchased from Sigma Aldridge (Saint Louis, MO). 1536-well plates were purchased from Greiner (cat # 789888).

NIMP may cause skin irritation, serious eye irritation and respiratory irritation. Alchem protocols consistent with safety and biosurety policies as well as EPA and Biohazardous material requirements were followed while working with NIMP.

Stocks of the reagents were made as follows: Amplex Red was made in DMSO at 5 mg/ml. Aliquots were frozen at -20°C and stored no longer than 2 weeks at a time. A stock of 5X assay buffer (250mM Tris, pH8) was filtered through a 0.2µm filter and stored at 4°C. On the day of the assay, the 5X buffer was diluted with sterile ultrapure water to 1X (50mM Tris, pH 8) and warmed to room temperature before beginning the assay. A stock of horseradish peroxidase was made in cold 1X assay buffer at 200U/ml. Aliquots were frozen at -80°C. A stock of choline oxidase was made in cold 1X assay buffer at 20U/ml and aliquots were frozen at -80°C. A stock of 100mM acetylcholine chloride was made in sterile ultrapure water, aliquoted and stored at -20°C. A stock of acetylcholinesterase (167U/ml) was prepared in cold 1X assay buffer and aliquots were stored at -80°C. NIMP was made fresh in DMSO for every set of experiments.

Equipment used for assay development included: a Labcyte/Beckman Coulter Echo 555 Acoustic transfer system (San Jose CA), a Beckman Coulter BioRAPTR FRD (Brea, CA), a Perkin Elmer Envision multi-mode plate reader (Waltham, MA) and an Eppendorf 5810 centrifuge (Hamburg, Germany).

### Basic Assay Protocol

NIMP is dispensed to a Greiner 1536 well plate using a Labcyte Echo 555 Acoustic transfer system. AChE is then dispensed to the plate using a Beckman Coulter BioRAPTR FRD. The plate is centrifuged in an Eppendorf 5810 centrifuge. Test compounds and control reactivators are dispensed to the plate with a Labcyte Echo 555 dispenser. The substrate mixture is dispensed to the plate using a Beckman Coulter BioRAPTR FRD and the plate is centrifuged with an Eppendorf 5810 centrifuge. The plate is then incubated at room temperature in the dark. Finally, the plate is read on a Perkin Elmer Envision multi-mode plate reader (Ex 555 and Em 590). Specific conditions will be presented in the results section.

### Data Analysis

Assay development data was analyzed using GraphPad Prism (GraphPad Software, Boston MA). The formula for percent reactivation is:

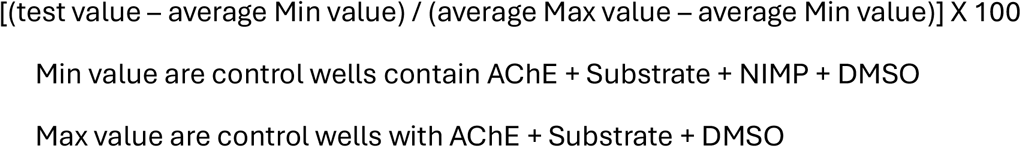

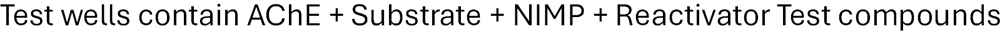

It should be emphasized that all wells contain the same percentage of DMSO. Reactivators are diluted in DMSO. NIMP is diluted in DMSO. Thus, when NIMP is added to the plates, 10nl of DMSO is added to the Max control wells. When reactivators are added to the plate, 10nl of DMSO is added to the Min control wells and Max control wells. When whole plates were run, the percent reactivation, Z score values and IC50 values were determined using CBIS software (ChemInnovation Software, Inc., San Diego, CA).

### Z’ score calculation for quality control

A statistical parameter has been developed which helps to determine if an assay is robust. This is known as the Z-factor or Z’. The Z-factor is established from four parameters: the means (μ) and standard deviations (σ) of both the positive (p) and negative (n) controls (μp, σp, and μn, σn) [18]. Z-factor is computed as:

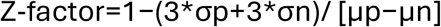

Z’ factor between 0.5 and 1.0 has been shown to represent a robust and reliable assay.

## Results

### Optimization and Miniaturization of the AChE Reactivation Assay Acetylcholinesterase 1536 Well Assay Development

Before initiating the development of the reactivator assay, we had previously developed an acetylcholinesterase inhibitor assay using the Amplex Red coupled assay. The assay conditions for the inhibitor assay were incorporated into the reactivator assay. The following processes and conditions were tested during the development of the 1536 well acetylcholinesterase inhibitor assay:

#### Titration and Time Course of Acetylcholinesterase

The amount of acetylcholinesterase used in the assay was determined with the goal to use as little enzyme as possible but with a quantity that would provide a robust and reproducible signal. A range of AChE concentrations from 0.025 to 0.25 units/ml was tested in the assay. It was determined that 0.025 U/ml was a suitable concentration for the assay (see Figure 2A).

**Figure 2:**
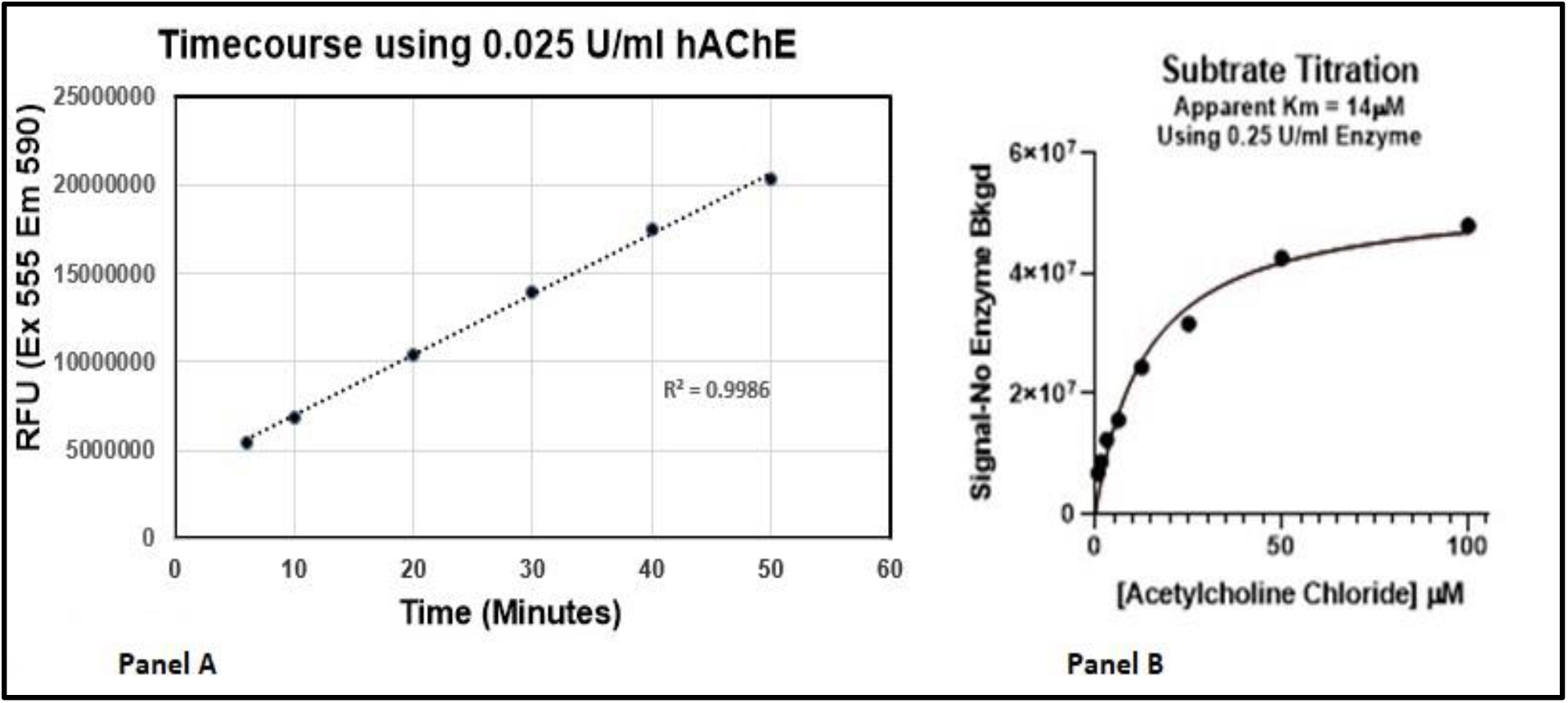
Acetylcholinesterase Time Course and Acetylcholine Titration. Panel A shows the timecourse using 0.025 U/ml of human AChE in a 4 µl assay. 2 µl of enzyme mixture (2 U/ml horseradish peroxidase, 0.2 U/ml choline oxidase and various concentrations of AChE in 50mM Tris pH 8.0) was added to the wells. 2 µl of substrate/Amplex Red mixture (30 µM acetylcholine chloride and 400 µM Amplex Red in 50 mM Tris pH 8.0) was added to the wells. The plate was incubated in the dark at room temperature and the fluorescence was read at various time points at ex 555 and em 590. Panel B shows the acetylcholine chloride titration in the 4 µl assay with conditions described in panel A details. In this assay 0.25 U/ml of hAChE was used.

#### Acetylcholine titration

For a biochemical assay, it is important to determine the Km for the substrate (assuming Michaelis–Menten kinetics; Km is the concentration of substrate which permits the enzyme to achieve half Vmax). Typically, the enzyme assay is run with the substrate at the Km. Because this assay is coupled, using 3 separate enzymes (AChE, choline oxidase and HRP) we will only be able to calculate the apparent Km for acetylcholine. Horseradish peroxidase and choline oxidase are not limited, and the concentrations are set at concentrations used in commercial kits for this Amplex Red coupled assay (final assay concentrations are 1 U/ml Horseradish Peroxidase and 0.1 U/ml choline oxidase). The assay is run at a fixed AChE concentration (0.25 U/ml) while varying the concentration of acetylcholine. It was determined that the apparent Km for acetylcholine in the assay is 14 μM. Data is shown in Panel B of Figure 2.

### Acetylcholinesterase Reactivation Assay Development

#### Initial data generated for HI-6 and 2-PAM

As previously stated, the initial assay conditions for the reactivation assay were based on the acetylcholinesterase inhibition assay. Initially, a 4 µl assay which followed the following steps was attempted: First, dilutions of NIMP were prepared in DMSO. Then, either 10 nl of the NIMP dilutions or 10 nl of DMSO for the positive controls were transferred to a Greiner 1536 well assay plate using a Labcyte/Beckman Coulter Echo 555 acoustic dispenser. Then, 2 µl of AChE (Final Assay Concentration (FAC) 0.025 U/ml) was dispensed using a Beckman Coulter BioRAPTR FRD. Next, the plate was centrifuged at 1000 rpm for 1 minute and incubated at room temperature for 30 minutes. During the incubation, dilutions of the reactivators (2-PAM and HI-6) were made in DMSO and transferred to an ECHO qualified 1536 well plate (in triplicate). After the 30-minute incubation, 10 nl of the reactivators was transferred from the ECHO source plate to the assay plate using a Labcyte/Beckman Coulter Echo 555 acoustic dispenser. Immediately, 2 µl of substrate mixture was added (final concentrations were 15 µM acetylcholine chloride, 200 µM Amplex Red, 0.1 U/ml Choline Oxidase and 1 U/ml horseradish peroxidase). The plate was centrifuged at 1000 rpm for 1 minute and then read on a Perkin Elmer Envision (excitation 555 and emission 590) every 10 minutes. Very weak activity was observed by 2-PAM and approximately 50% reactivation with HI-6. Because 2-PAM was a weak reactivator compared to HI-6, HI-6 was used for subsequent assay development experiments. Initial Data is shown in Supplemental Figures 1 and 2. This data was generated with FAC of NIMP set at 0.059 nM.

#### DMSO Tolerance

For the acetylcholinesterase inhibitor assay, the dimethylsulfoxide (DMSO) exposure is not only from the test compounds but also the NIMP. When titrated in the inhibitor assay, DMSO had a slight effect above 0.25% (Supplement Figure 3). However, in the AChE reactivator assay, the enzyme is first exposed to NIMP which is solubilized in DMSO and when the reactivator test compounds are added, these are also solubilized in DMSO which doubles the DMSO concentration. Plus, we believe there is extra stress on the enzyme from the phosphorylation by NIMP which might reduce the enzyme’s tolerance to the DMSO. With these considerations in mind, it was decided to measure the AChE activity in a 4 µl and an 8 µl assay. To do this, the enzyme was exposed to varying concentrations of DMSO delivered to the plates with the Labcyte/Beckman Coulter Echo 555 acoustic dispenser. Choline oxidase and horseradish peroxide are added with the substrate and Amplex Red (FAC:15 µM acetylcholine chloride, 1U/ml Horseradish Peroxidase, 0.1 U/ml choline oxidase, 200 µM Amplex Red in 50 mM Tris, pH8.0). This method of addition was used for all subsequent experiments. The data shown in Figure 3 demonstrates that the enzyme functions better in the 8µl assay with the optimal DMSO concentration no higher than 0.25%. Thus, it was decided that the assay would be an 8µl assay using 10 nl of NIMP and 10 nl of test compound (reactivators).

**Figure 3:**
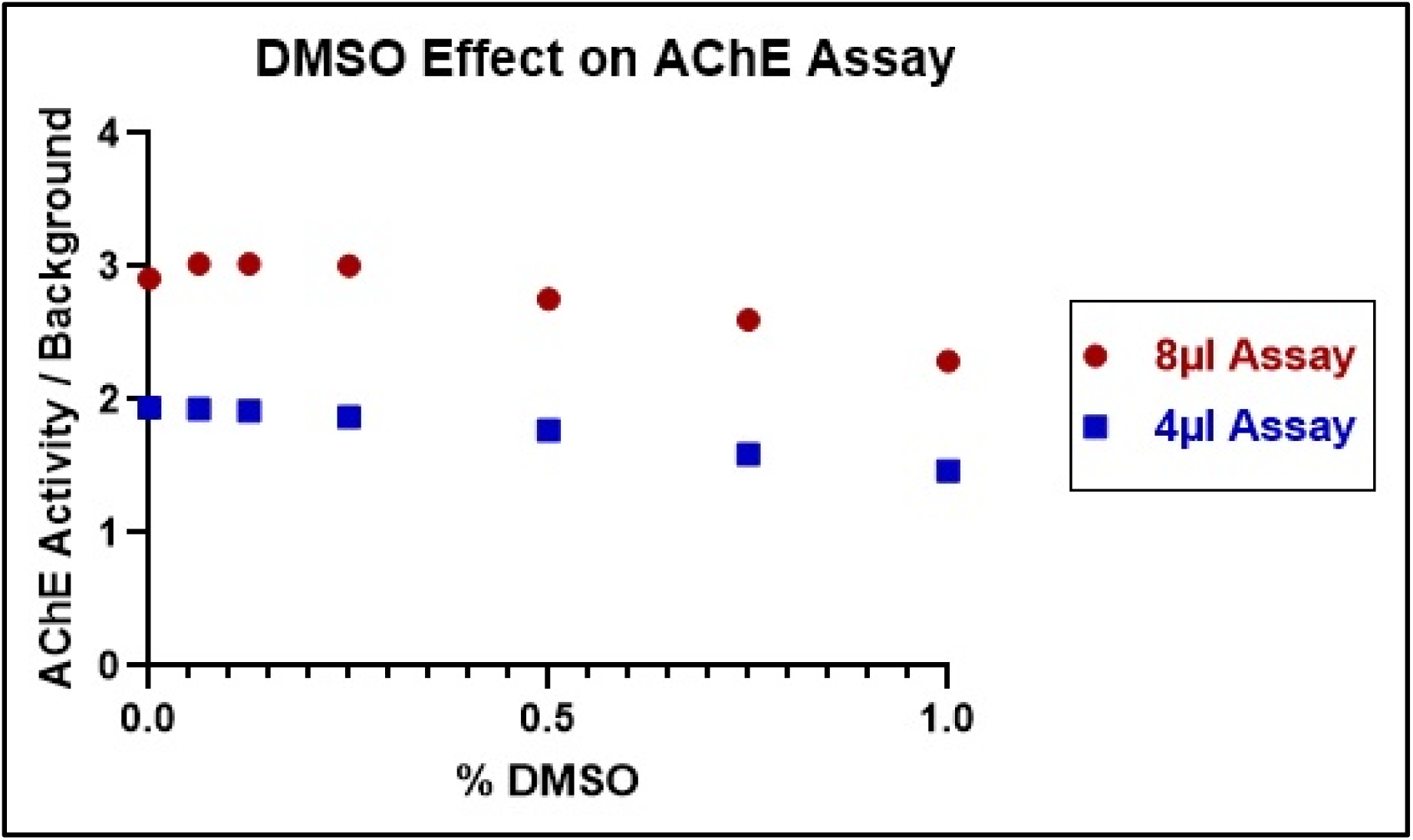
DMSO Effect on AChE Activity. Varying amounts of DMSO was dispensed to 2 assay plates using the ECHO 555 acoustic dispenser. Then, 2 µl of AChE enzyme (final concentration 0.025 U/ml) was dispensed to the top 16 rows and 2 µl of buffer was dispensed to the bottom 16 rows of plate 1. Likewise, 4 µl of enzyme was dispensed to the top 16 rows and 4 µl of buffer was dispensed to the bottom 16 rows of plate 2. The two plates were centrifuged for 1 minute at 1000 rpms. Then 2 µl substrate was dispensed to all wells of plate 1 and 4 µl was dispensed to all wells of plate 2. The plates were centrifuged again and incubated for 30 minutes at room temperature in the dark. Then the plates were read on the Envision at Ex 555 and Em 590.

#### NIMP Titration in 8 µl AChE Reactivation Assay

Once it was confirmed that the AChE activity improved in the 8 µl assay, the next step was to test NIMP in the 8 µl assay. Dilutions of NIMP were made in DMSO and 10 nl of each dilution was transferred to the assay plate using an ECHO 555 acoustic dispenser and 4 µl AChE was dispensed using a BioRAPTR FRD. The plate was centrifuged for 1 minute at 1000 rpm and 4 µl substrate mixture (FAC:15 µM acetylcholine chloride, 1 U/ml Horseradish Peroxidase, 0.1 U/ml choline oxidase, 200 µM Amplex Red in 50 mM Tris, pH 8.0) was added to the plate. The plate was centrifuged and incubated in the dark at room temperature for 30 minutes. The plate was read on a Perkin Elmer Envision with Ex 555 and Em 590. Because it is easy to completely kill the enzyme with NIMP, we wanted to work at a NIMP concentration just above baseline (EC75). Based on NIMP dose response curves, the concentration for NIMP that gives EC75 was determined to be 0.09 nM (data shown in figure 4).

**Figure 4:**
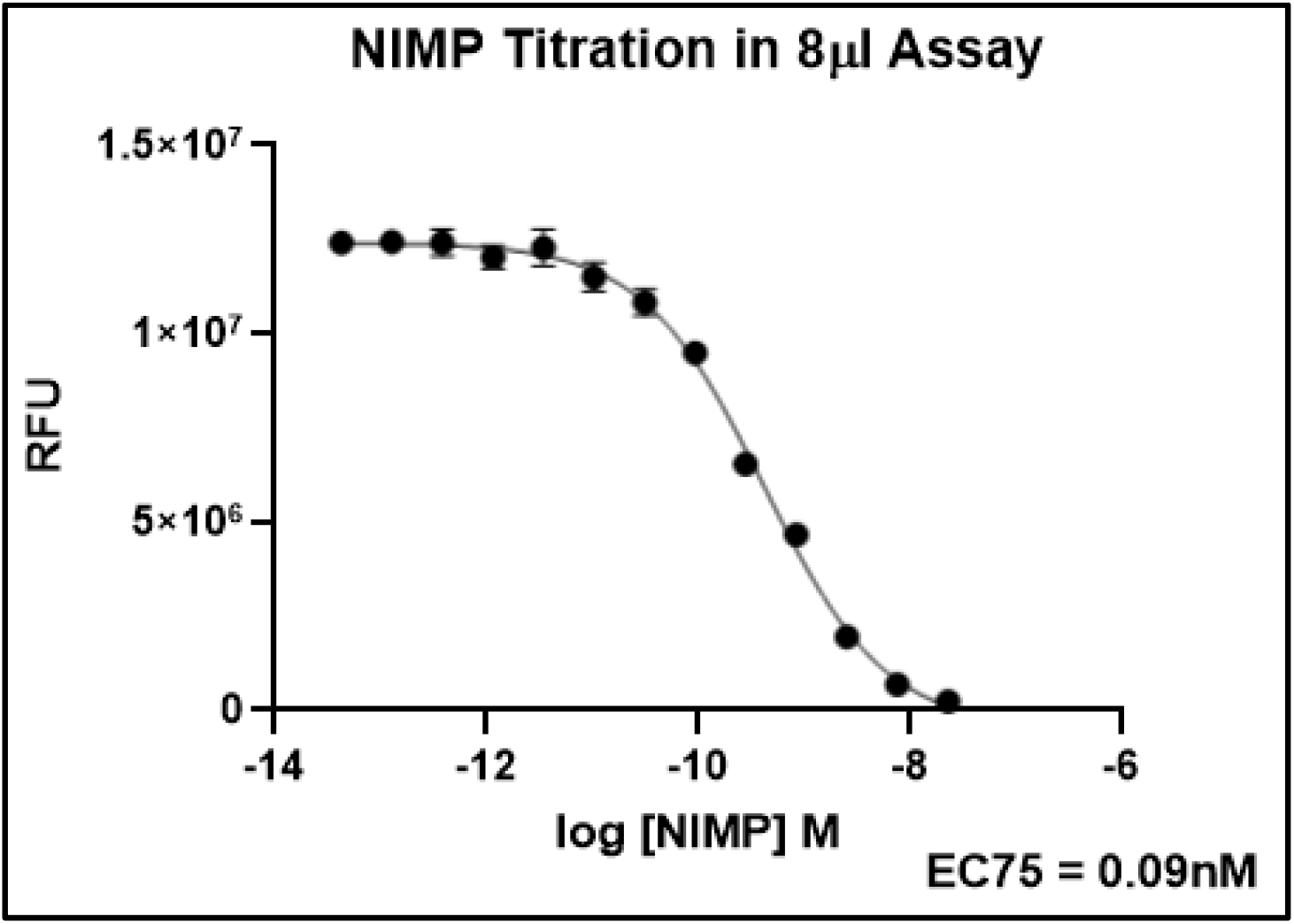
EC75 of NIMP in the 8µl AChE Reactivation Assay. NIMP was diluted in DMSO and transferred to an assay plate using the ECHO 555 acoustic dispenser. Then, 4µl of AChE enzyme (final concentration 0.025U/ml) was dispensed to wells. The plate was centrifuged for 1 minute at 1000 rpms. Next, 4µl substrate mixture (FAC:15 µM acetylcholine chloride, 1 U/ml Horseradish Peroxidase, 0.1 U/ml choline oxidase, 200 µM Amplex Red in 50 mM Tris, pH 8.0) was dispensed to the plate. The plate was centrifuged and incubated for 30 minutes at room temperature in the dark and read on the Envision at Ex 555 and Em 590. Data was fitted with a four-point logistical model in GraphPad Prism.

#### HI-6 Titration

The control reactivator (HI-6) was titrated to determine the best concentration for the assay. For this experiment, NIMP was dispensed at a final concentration of 0.09nM (10 nl) using the ECHO 555. Then 4 µl of AChE (final concentration 0.025 U/ml) was dispensed with a Beckman Coulter BioRAPTR The plate was centrifuged at 1000 rpm for 1minute and immediately 10 nl of HI-6 diluted in DMSO at various final concentrations was dispensed to the plate using a Labcyte Echo 555. The plate was centrifuged again at 1000 rpm for 1 minute and incubated for 15 minutes at room temperature. Then 4 µl of the substrate mixture (FAC:15 µM acetylcholine chloride, 1 U/ml Horseradish Peroxidase, 0.1 U/ml choline oxidase, 200 µM Amplex Red in 50 mM Tris, pH 8.0) was dispensed to the plate. The plate was centrifuged at 1000 rpm for 1 minute, then read on a Perkin Elmer Envision every 10 minutes for a total of 90 minutes. The data (shown in figure 5) indicates that 6.25 µM HI-6 is the optimal concentration for the assay because 12.5 µM HI-6 quenches the signal. HI-6 at 6.25 µM yields roughly 73% reactivation at 90 minutes in this experiment.

**Figure 5:**
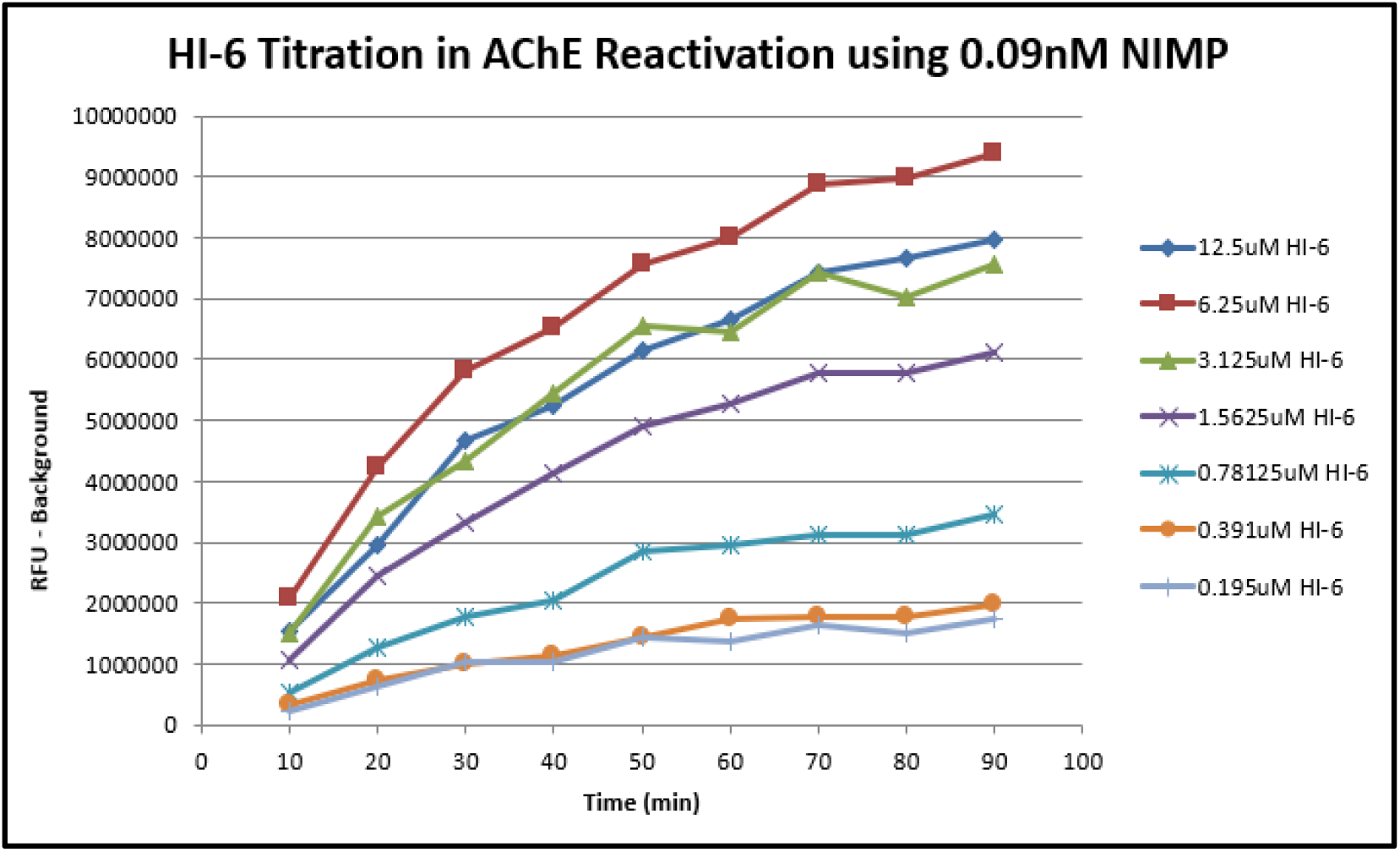
HI-6 Titration in AChE Reactivation Assay: Dispensed NIMP (FAC 0.09 nM) to the assay plate using a Labcyte Echo 555. Then, dispensed 4 µl of AChE enzyme using a Beckman Coulter BioRAPTR. The plate was centrifuged at 1000rpm for 1 minute and immediately 10 nl of HI-6 at various concentrations was dispensed to the plate using a Labcyte Echo 555. The plate was centrifuged again at 1000 rpm for 1 minute and incubated 15 minutes at room temperature. Then 4 µl substrate mixture (FAC:15 µM acetylcholine chloride, 1 U/ml Horseradish Peroxidase, 0.1 U/ml choline oxidase, 200 µM Amplex Red in 50 mM Tris, pH 8.0) was added to the plate. The plate was centrifuged at 1000 rpm for 1 minute and then read on a Perkin Elmer Envision every 10 minutes.

#### Final Optimization of the AChE Reactivation Assay

When testing compounds in the assay, it was discovered that incorporating a titration of the NIMP before each experiment proved beneficial. The goal was to refine the NIMP concentration to get 50% reactivation by 6.25 µM HI-6 in 30 minutes. In this way variation caused by NIMP, which is a compound that is easily oxidized and will readily lose activity upon oxidation, could be controlled. With that said, generally the deviation from 0.09 nM NIMP is only a modest shift from experiment to experiment. A time course for the assay was also incorporated, taking readings every 30 minutes up to 5 hours. By doing this, fluorescent artifacts are identified because their fluorescence will not increase with time. Also, fluorescent quenchers have a diminished effect at 5 hours as the enzyme converts the substrate. In every experiment we include a dose response of the two control compounds (HI-6 and 2-PAM). At the 5-hour time point, the EC50 is calculated for each compound using the CBIS software. Shown in Figure 6 are representative curves of both 2-PAM and HI-6 in our assay at the 5-hour time point and the average EC50 values for all our experiments to date.

**Figure 6:**
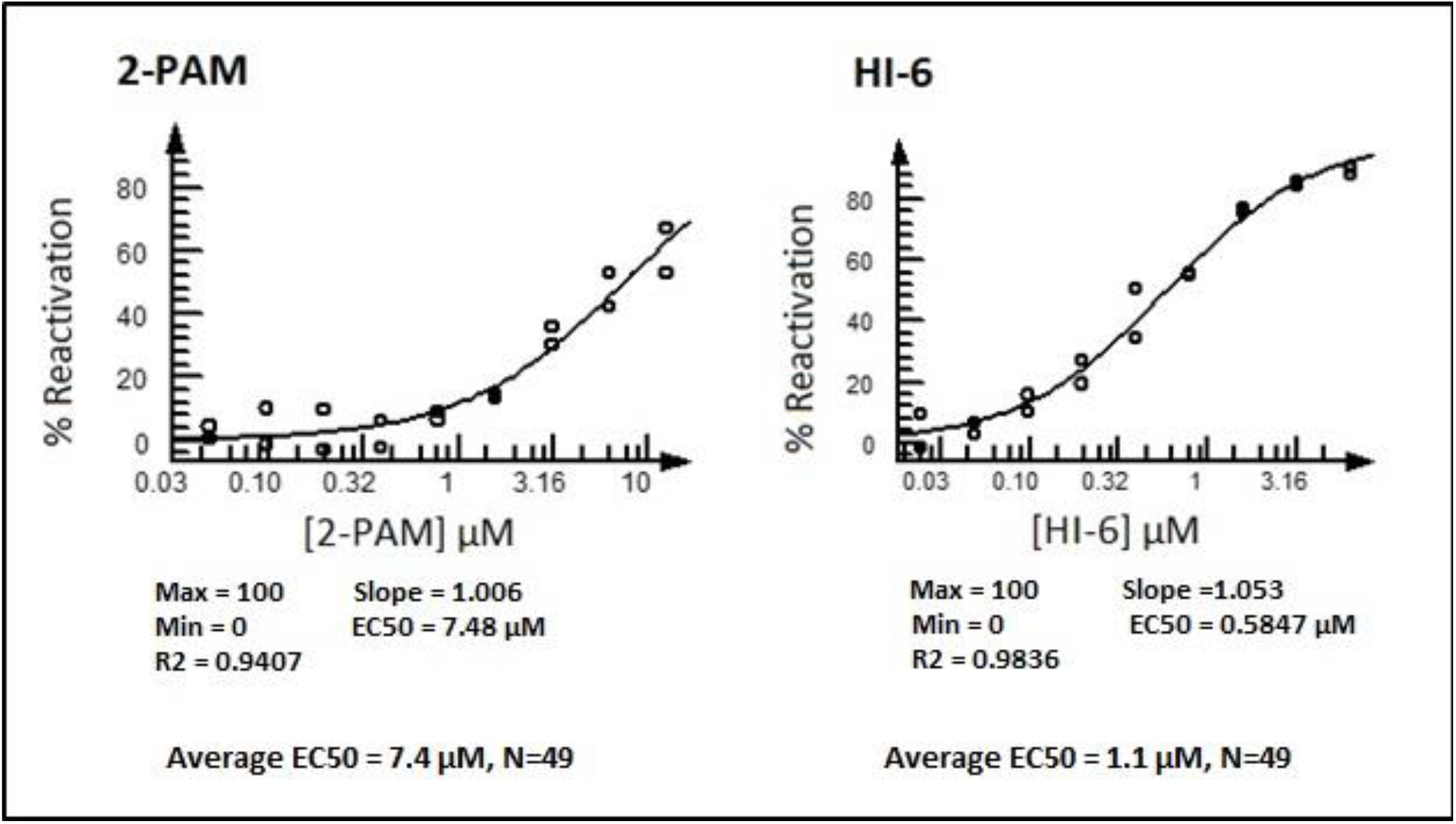
Dose Responses of 2-PAM and HI-6 in AChE Reactivation Assay. The 5-hour data was analyzed using the CBIS software. The average values shown at the bottom of the figure were from 49 individual experiments.

### Final Acetylcholinesterase Reactivation Assay Protocol

Based on the initial developmental studies described above, the following protocol has been developed.

1. Titrate the NIMP so that the concentration yields 50% Reactivation by 6.25 µM HI-6 at 30 minutes.
2. Dispense 10 nl of NIMP or DMSO to a 1536-well Greiner assay plate. The NIMP is dispensed into all columns except for columns 3 and 4. DMSO is dispensed into columns 3 and 4 which are the 100% enzyme activity control wells.
3. Dispense 4 µl of human acetylcholinesterase (FAC are 0.025 U/ml) to all wells of the plate using the BioRAPTR FRD.
4. Centrifuge the plate for 1 minute at 1000 rpm in an Eppendorf centrifuge.
5. Immediately dispense 10 nl test and control reactivators to the plate (DRCs of control reactivators 2-PAM and HI-6 are included). Dispense 10 nl of DMSO into columns 1 which is the 0% reactivation controls and columns 3 and 4 (again, these are 100% enzyme activity control wells). Dispense 10 nl of HI-6 to column 2 (FAC 6.25 µM) for the reactivation control.
6. Dispense 4µl of substrate mixture (FAC:15 µM acetylcholine chloride, 1 U/ml Horseradish Peroxidase, 0.1 U/ml choline oxidase, 200µM Amplex Red in 50 mM Tris, pH 8.0) to all wells of the assay plate using a BioRAPTR FRD.
7. Centrifuge the plate for 1 minute at 1000 rpm in an Eppendorf centrifuge.
8. Read the plate on a Perkin Elmer Envision (Ex 555 and Em 590). For screening compounds at a single concentration, two readings are taken at 30 minutes and 5 hours. If the precent reactivation does not change, the compound is a fluorescent artifact. For dose response curves, readings are taken every 30 minutes to evaluate the speed which the reactivation occurs. The EC50 curve is generated using the 5-hour time point.

Using this protocol good plate statistics were consistently obtained, with an average Z value of 0.72 and an average signal window of 5.6-fold. Average CV is 7.4%.

### Validation with Compounds from an *In Silico* Virtual Screen

To validate the AChE reactivation assay, we conducted an *in silico* screen of 20,000 oxime compounds from the Chemspace oxime small molecule library using an oxime reactivator docking model to rank candidates for assay validation. The human homology oxime reactivator docking model was developed in collaboration with Molecular Forecaster Inc. The model, which will be described in detail elsewhere, was based on the mouse sarin-inactivated AChE crystal structure (PDB ID: 2WHP) complexed with the oxime reactivator HI-6. The 2WHP structure is unique in having the oxime oxygen function of the oxime in the correct directional alignment to mediate reactivation by nucleophilic attack on the phosphorus atom of the phosphorylated active site AChE-OP serine residue (see Figure 7 below). The reactivator docking model was optimized from the X-ray structure using a molecular dynamics approach, simulating the oxime attack on phosphorus. A set of 42 oximes were selected from the 20,000 oxime compound library screen based on high docking score ranking, chemical diversity and commercial availability.

**Figure 7:**
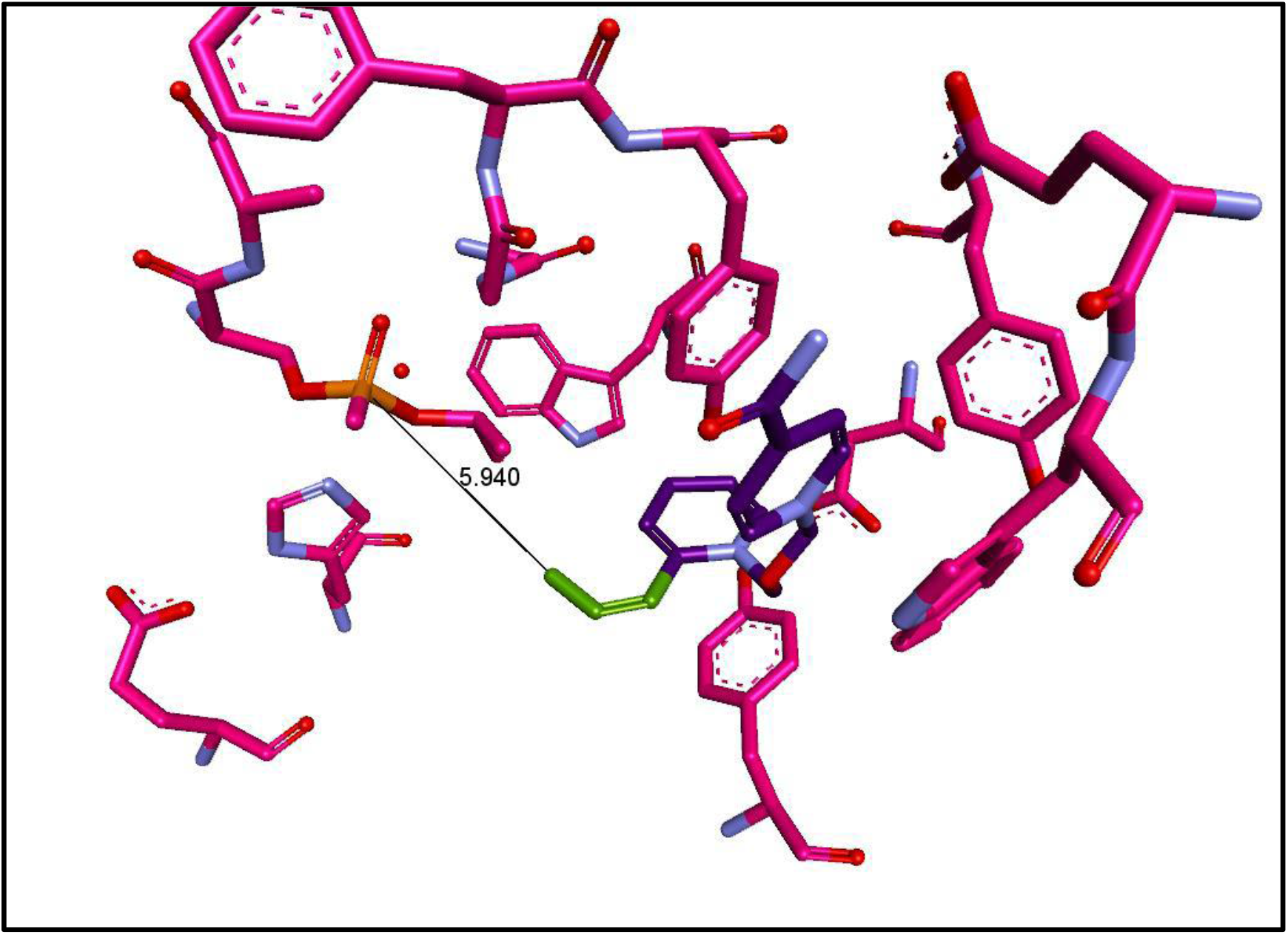
PDB Crystal Structure 2-WHP, Mouse AChE-OP derived from Sarin in “productive” Complex with HI-6 Reactivator. The HI-6 oxime oxygen (green) is oriented toward the phosphorus of AChE-OP at a distance of 5.9 angstrom.

From the NIMP assay screen of the 42 in silico selected compounds, hit oxime reactivator Compound **8** was identified with an average EC50 = 4.1 µM (N=3). The data for this validation screen of the 42 compounds is shown in Figure 8. Figure 9 shows the dose curves for Compound **8** in the 3 assays. Chemical lead optimization from Compound **8** resulted in several novel oxime reactivator compounds with greater potency. One of these novel oxime reactivators, Compound **127** is more potent and faster acting than the current gold standard reactivator HI-6 in our assay.

**Figure 8:**
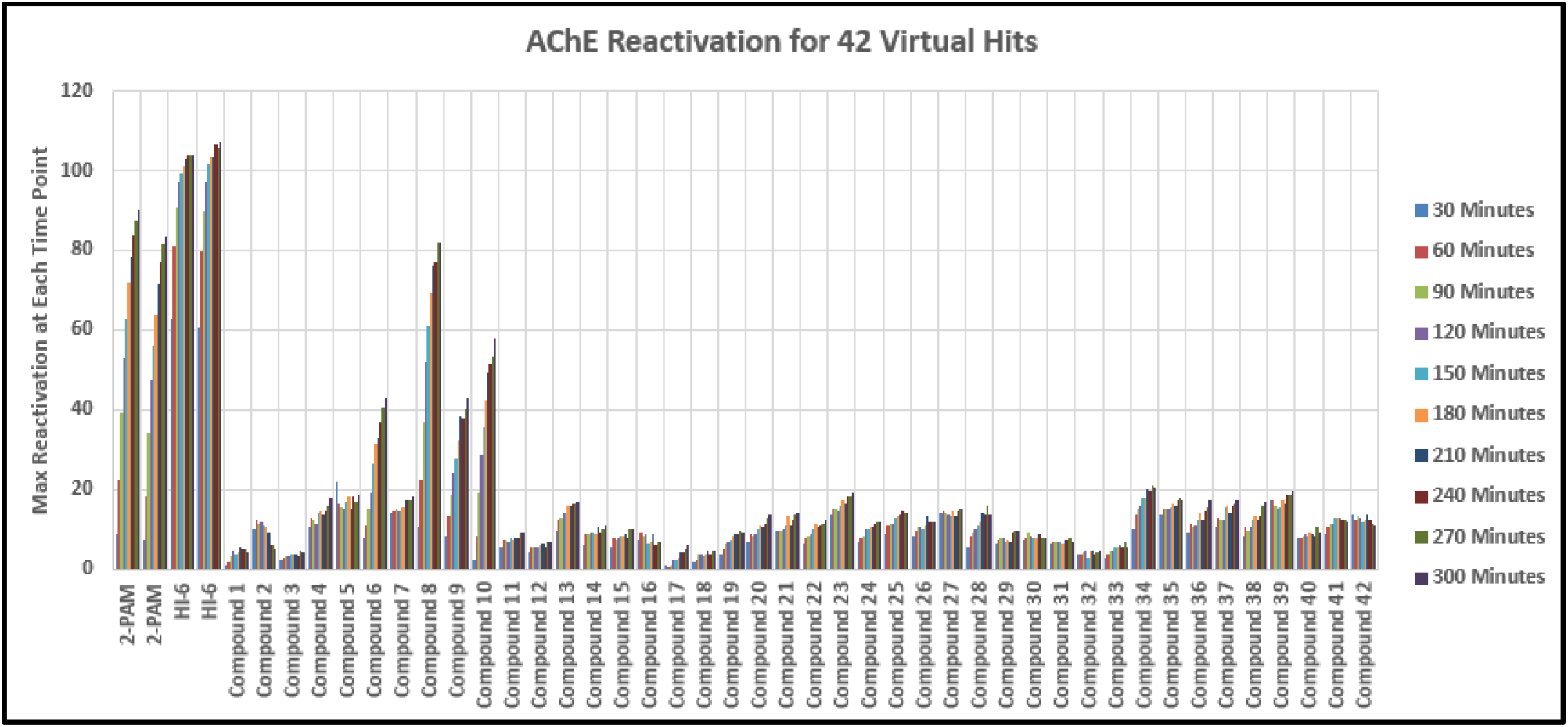
Results of Validation Screen for the AChE Reactivation Assay. An *in silico* virtual screen was performed using a library of 20,000 oxime small molecules. Forty-two compounds were tested in this experiment in dose response using the protocol described in “Final Acetylcholinesterase Reactivation Assay Protocol” section.

**Figure 9:**
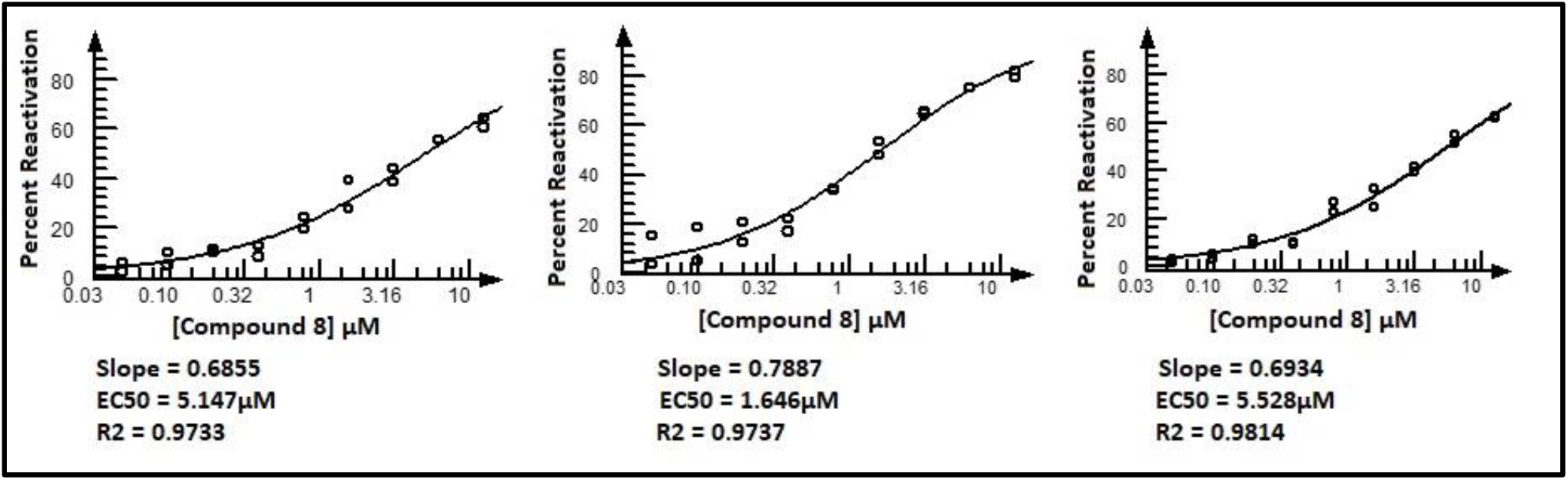
Dose Response Curves for Compound 8 in the AChE Reactivation Assay. Dispensed NIMP (FAC 0.09nM) to the assay plate using a Labcyte Echo 555. Then, dispensed 4 µl of AChE enzyme using a Beckman Coulter BioRAPTR. The plate was centrifuged at 1000rpm for 1 minute and immediately 10nl of test compounds along with control reactivators were dispensed into the assay plate using a Labcyte Echo 555. Then 4 µl substrate mixture (FAC:15 µM acetylcholine chloride, 1 U/ml Horseradish Peroxidase, 0.1 U/ml choline oxidase, 200 µM Amplex Red in 50 mM Tris, pH 8.0) was added to the plate using the BioRAPTR. The plate was centrifuged at 1000 rpm for 1 minute, incubated in the dark and read on a Perkin Elmer Envision (Ex 555 and Em 590). The data shown in this figure is from the 5-hour time point. The data was analyzed using CBIS. The curves shown here are from 3 different experiments on 3 different days.

## Conclusion

Our primary goal for developing this high-throughput AChE reactivation assay was to discover novel reactivators of organophosphate poisoned acetylcholinesterase (AChE-OP) with improved broad spectrum against OP nerve agents vs 2-PAM and HI-6. The assay described in this paper allows quick analysis of test compounds in a convenient and a safe 1536-well format. The assay can be used for either a high-throughput single concentration screen using 2 time points or for dose responses as described in this paper. Here we validated the assay with 42 oxime reactivator test compounds selected from a virtual screen along with 2 known reactivators, namely 2-PAM and HI-6. One of the compounds from the validation screen (Compound **8**) had an EC50 = 4.1 µM (N=3). Further analog synthesis and structure activity optimization from Compound **8** led our team to Compound **127** which demonstrated rapid reactivation of AChE-OP in our NIMP assay with potency equivalent to HI-6. The translational relevance of the NIMP assay reactivator activity of Compound **127** from testing across a panel of OP nerve agents including sarin will be detailed in a subsequent report. Thus, this assay provides a critically needed tool to advance the identification of novel AChE reactivators.

