## Supplement data for "Development of a High-throughput Assay to Identify Novel Reactivators of Acetylcholinesterase"

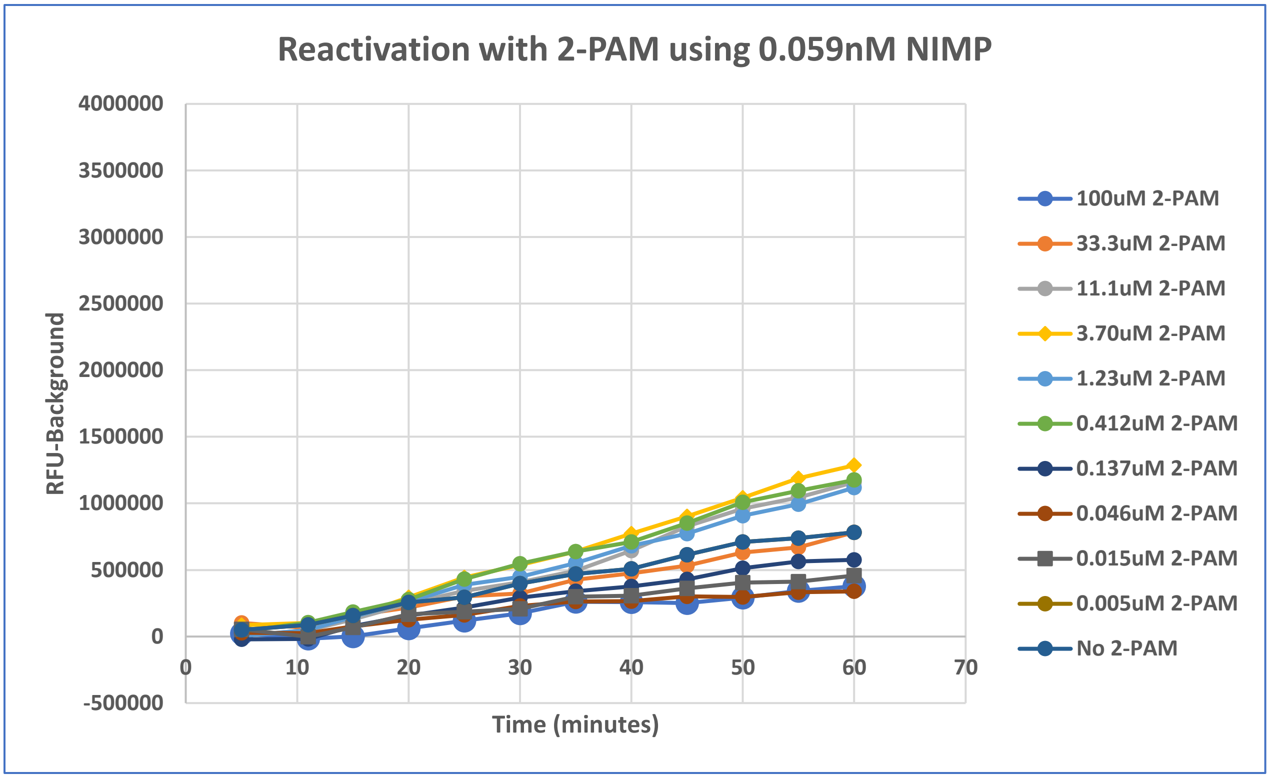


**Supplement Figure 1: Titration of 2-PAM using 0.59nM NIMP.** NIMP diluted in DMSO was transferred to the assay plate using a Labcyte/Beckman Coulter Echo 555 acoustic dispenser (FAC 0.059nM, 10 nl). Then, 2 µl of AChE was dispensed using a Beckman Coulter BioRAPTR FRD. The plate was centrifuged at 1000 rpm for 1 minute. After a 30-minute incubation at RT, 2-PAM (diluted in DMSO at various concentrations) was dispensed using the Echo (10 nl of each concentration) and 2 µl substrate was added. The plate was centrifuged at 1000 rpm for 1 minute and then read on the Envision every 5 minutes.

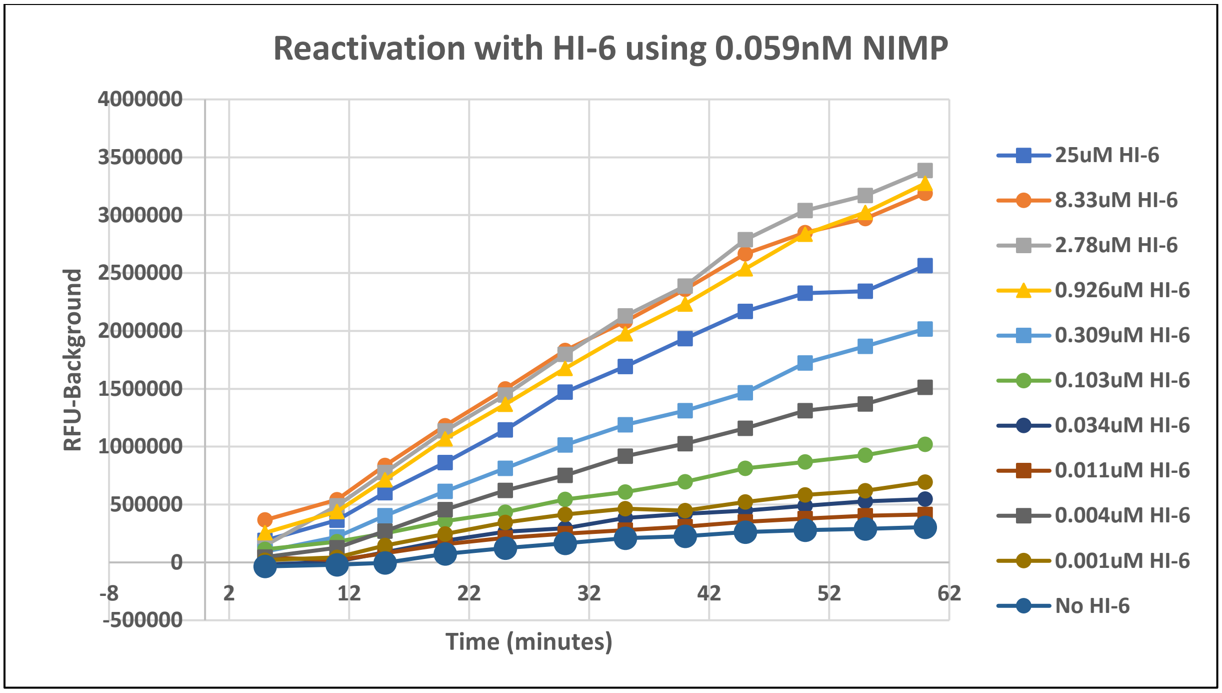


**Supplement Figure 2: Titration of HI-6 using 0.59nM NIMP.** NIMP was transferred to the assay plate using the Echo 555 acoustic dispenser (FAC 0.059nM, 10 nl). Then, 2 µl of AChE was dispensed using the BioRAPTR. The plate was centrifuged at 1000 rpm for 1 minute. After a 30-minute incubation at RT, 2-PAM (diluted in DMSO at various concentrations) was dispensed to the assay plate. The plate was centrifuged and 2 µl substrate mixture was added to all wells. The plate was centrifuged at 1000 rpm for 1 minute and then read on the Envision every 5 minutes.

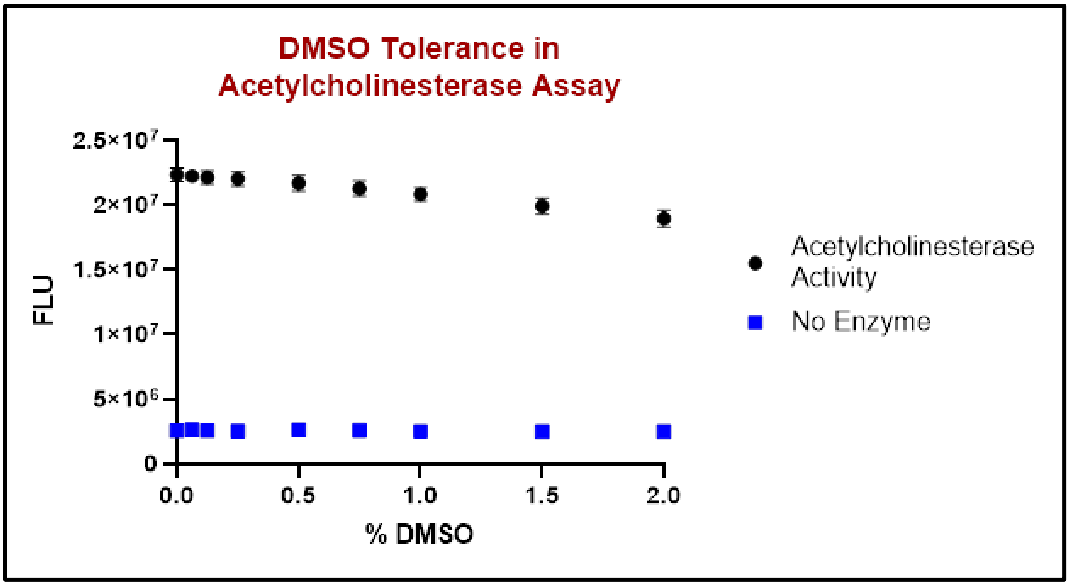


**Supplement Figure 3: DMSO Tolerance**. Various amounts of DMSO are added to the wells. 2 µl of enzyme mixture (1 U/ml horseradish peroxidase, 0.1 U/ml choline oxidase and 0.25 U/ml AChE in 50 mM Tris pH 8.0) were added to the wells. 2 µl of substrate/Amplex Red mixture (15 µM acetylcholine chloride and 200 µM Amplex Red in 50 mM Tris pH 8.0) were added to the wells. The plate was incubated in the dark at room temperature for 35 min, then fluorescence was read at ex 555 and em 590.
